# Automahout: an autonomous image analysis system for cell tracking

**DOI:** 10.64898/2026.11.04.764293

**Authors:** Ko Sugawara, Shuichi Onami

## Abstract

Cell tracking in time-lapse images is complex image analysis that involves detection, linking and proofreading, often requiring the algorithm to be tuned for every new dataset. We present Automahout, an autonomous agent for the Mastodon/ELEPHANT platform that carries out these operations. It looks at the data and tries existing models; where none fits, it annotates reference cells, trains a detector from those sparse annotations, reviews its output against them and feeds the corrections back until its own stopping criteria are met. It then links, checks the lineage and moves on. On three public datasets, a 2D fluorescence culture, a 2D phase-contrast culture and a 3D embryo, it produced lineages without human input and reached TRA scores of 0.95–0.96.

## Introduction

Live imaging follows cells over hours to days, from cultured cells to developing tissues and whole organisms. Questions such as when a cell divides, where its daughters go and how a lineage is apportioned require complete and accurate lineages. Achieving this is complex image analysis. Cells must be detected in every frame and linked across frames, and the result must be reviewed and corrected where detection or linking went wrong. Detection can use conventional rule-based algorithms with tuned parameters or trained models, and generalist pretrained models such as StarDist [1, 2], micro-SAM [3] and Cellpose-SAM [4] now detect a wide range of cells without retraining. However, none works on every dataset, especially in 3D, where cells are crowded and image quality changes with depth. A method that works under one set of conditions may therefore fail when the specimen, label, magnification or stage changes. In practice, a person repeats these operations for every new dataset, judging from the images what is wrong and how to fix it, until the result is good enough. Even as methods improve, this reliance on a person’s judgement remains.

ELEPHANT made these operations efficient for deep-learning-based tracking [5]. It trains detection and linking models from sparse annotations rather than densely labelled volumes, and it puts the person inside the training loop: a few annotated cells train a model, the person corrects its predictions in Mastodon [6], the corrections become the next round of training data, and accuracy rises quickly. What it did not change is who performs these operations. What to annotate, when to train, which predictions are wrong, how to fix them and when to stop are all judgements made by a person looking at the images, and they are made again for every dataset.

Large language models (LLMs) and vision-language models (VLMs) have made it possible to build agents that plan a task, call software tools and inspect what comes back [7, 8]. In bioimage analysis, such agents have begun to write analysis code, operate GUI image-analysis software such as ImageJ/Fiji and napari (Omega [9], Agentic-J [10], CopilotJ [11]) and answer questions about tools and run them on request (BioImage.IO Chatbot [12]). An agent of this kind could take the person’s place in tracking, provided it can also judge the result from the images, as a person does.

Here we present Automahout, an autonomous agent that carries out these operations. It works on the Mastodon/ELEPHANT platform, makes the same kinds of annotation a person makes there, and judges its own output from the images, without ground truth. We show it on three public datasets that call for different operations.

## Results

### Automahout carries out the operations of tracking

Automahout is a Mastodon plugin in Fiji [13]. A person opens a Mastodon project, starts Automahout from Mastodon’s Plugins menu and watches it work in Mastodon’s own views, where every annotation, detection and link it makes can be inspected and edited (Supplementary Fig. 4); the runs reported here ran the same plugin code as batch jobs from the command line, with no interaction at any point. It carries out the operations in the order a person does, using the same ELEPHANT server and the same kinds of annotation (Fig. 1a). Like a person, it starts by looking at the data, including the modality, the cell size and what a vision-language model sees in frames sampled across the whole movie. It then tries existing pretrained detectors and keeps one only if its detections agree with the image’s bright regions, on 2D fluorescence, or otherwise are consistent in space and time, as real cells are. It then works through the movie in time windows of 10 frames, finishing each before moving to the next, so that it can adapt its settings, and the model when it trains its own, as the cells and the images change over time. In each window it reads the cell size again from the image, so that it follows cells that shrink as an embryo develops (Fig. 1c). If no pretrained detector passed, it trains its own. It annotates reference cells on frames of the window, trains an ELEPHANT detector from those sparse annotations, runs it on the whole frame, and evaluates the result against the references as a person would (Fig. 1b): which cells it found, which it missed and which detections are on nothing. Missed cells and false detections go back as ELEPHANT’s own correction labels, and the model is trained again. Once detection in the window is settled, it links the detections over time and checks the lineage for what real lineages do not do. It looks for divisions that come too soon after the previous one or whose daughters do not move apart, for breaks, for tracks too short to be a cell and for one cell tracked twice, and corrects them. Before it moves to the next window, it trains its own detection model once more on the window’s finished result, with each cell’s shape fitted to the image, as a person does after proofreading.

**Figure 1.**
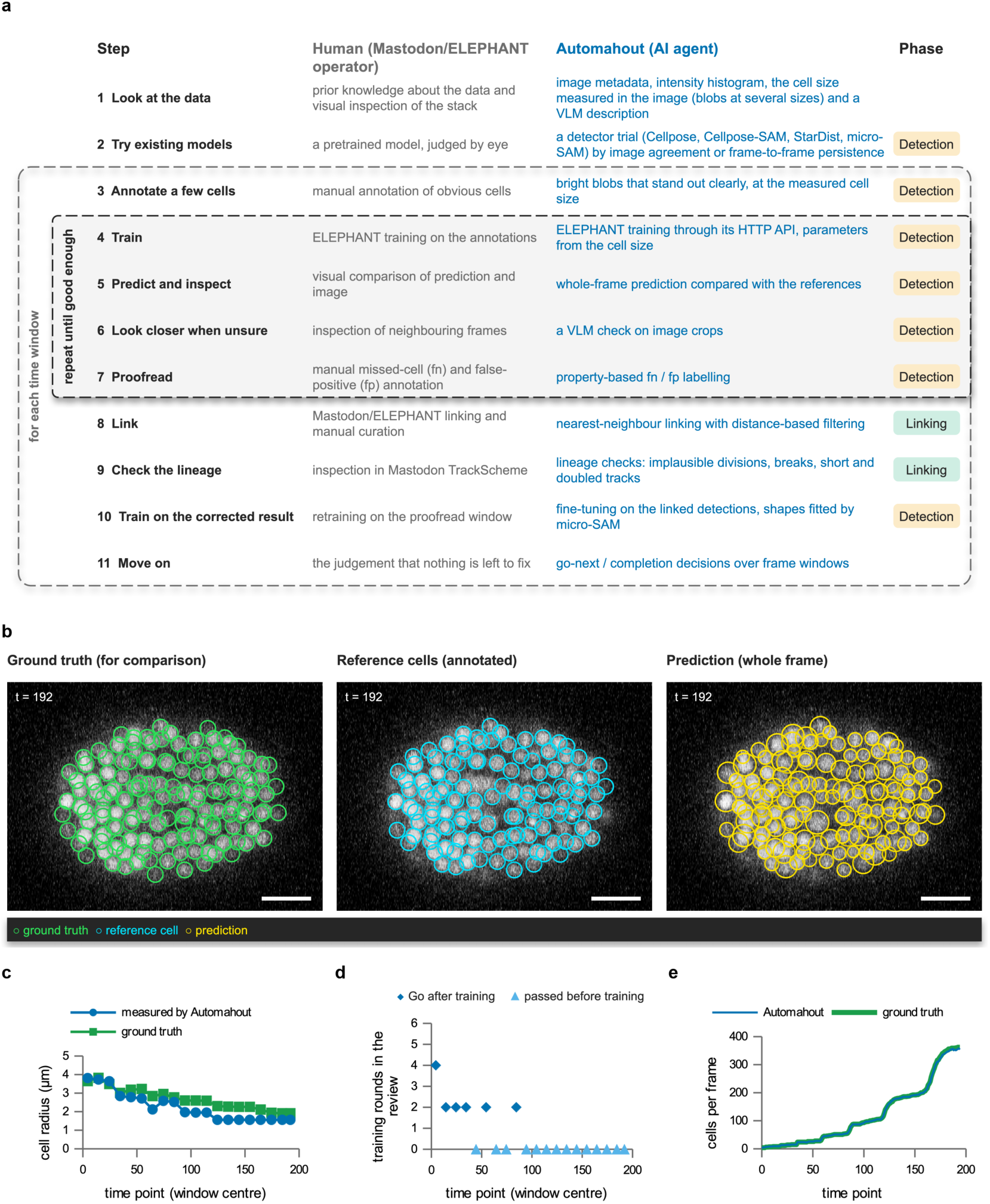
| Automahout carries out the operations of tracking. (**a**) The steps of tracking a new dataset, each with how a person carries it out in Mastodon/ELEPHANT and how Automahout does; the boxes mark the training loop, repeated until good enough, and the steps repeated for each time window; the last column gives each step’s phase, detection (2–7, 10) or linking (8–9). (**b**) One review round on the embryo (t = 192): the ground truth for comparison, the reference cells Automahout annotated, and the model’s whole-frame prediction (green, cyan and yellow circles). The references are a sample of the cells (115 of the 143 drawn): the confident ones. Maximum-intensity projection over z = 21.5–26.5 µm, with circles for cells centred within one cell radius (1.56 µm) of it, grayscale, display range 23–255 (1st–99.8th percentile); scale bar, 10 µm. (**c**) The cell radius Automahout measured per time window by scale selection, against the ground truth’s (the challenge’s silver truth, the only segmentation of every frame), as the embryo’s cells shrink. (**d**) Training rounds each window’s review needed: Go after training, or passed before any training. (**e**) Cells per frame, ground truth and Automahout. Training runs where the cell size changes most (c, d), as the embryo’s cells divide (e).

### Deciding when to stop, and looking closer when unsure

A person stops training when there is nothing left to fix. Automahout applies the same test in its review. It stops when, in two rounds running, the detector finds all but a few per cent of the references, makes almost no detections on nothing, and draws cells at about their size. Because a single good round can be luck, it waits for a second. If the review never gets there, Automahout keeps the best round rather than the last, as a person would undo a round of training that made things worse (Fig. 1d). A person who cannot tell whether something is a cell looks more closely. Automahout does the same when one reference keeps being missed while everything else is right. It shows that reference to the vision-language model and asks whether it is a nucleus (Supplementary Fig. 1); if not, the reference is dropped and the detector is no longer required to find it.

### Tracking three datasets without a person

We ran Automahout without human input on the first training sequence (01) of each of three Cell Tracking Challenge datasets [14, 15], where the ground truth is public, chosen because they call for different operations (Fig. 2, Table 1): a 2D fluorescence culture (Fluo-N2DL-HeLa), a dense 2D phase-contrast culture (PhC-C2DL-PSC), and a *C. elegans* embryo imaged in 3D (Fluo-N3DH-CE), which grows from 4 to 362 cells while the median radius of their nuclei shrinks from 4.3 to 1.9 µm. On the two cultures, a pretrained detector passed its check, and Automahout chose Cellpose-SAM [4] on both: on HeLa it agreed best with the image’s bright regions, and on PSC its detections persisted best from frame to frame, also in the dense late frames. Automahout used it as it is, then linked the detections and corrected the lineage (HeLa DET 0.954, TRA 0.954, BC(0) 0.83; PSC DET 0.956, TRA 0.954). When Cellpose-SAM was not yet among the candidates, the same check chose micro-SAM [3] on HeLa and Cellpose cyto3 [16] on PSC, and PSC scored DET 0.904 and TRA 0.893: adding a better model was enough for Automahout to take it up (Supplementary Table 1). A baseline that applies one pretrained detector as it is and links its masks with Trackastra [17] scored lower on both cultures (TRA 0.929 on HeLa, 0.900 on PSC) and failed on the embryo (TRA 0.202), where no pretrained detector found the cells. On the embryo no pretrained detector passed, and Automahout therefore carried out every operation itself. It trained an ELEPHANT detector from random weights on reference cells it had annotated, reviewed and corrected it window by window, and followed the cells as they shrank (Fig. 1c–e). Every window reached Go, most within a few rounds. The vision-language model was consulted once, at t = 5, and judged one doubtful reference not to be a nucleus, although it was a real cell; the reference was dropped (Supplementary Fig. 1). Over the frames with ground truth (t ≤ 194) Automahout reached DET 0.971 and TRA 0.965 (LNK 0.920, BC(0) 0.55). The errors gather where the embryo crowds, after t ≈ 160: summed over those frames, 252 cells were missed, and of the detections 248 were duplicates on one cell, 184 merged two cells and 26 lay on no cell (Supplementary Fig. 2). No question was put to a person in any of these runs (Supplementary Fig. 3); the runs took 31 min (HeLa), 49 min (PSC) and 4.5 h (embryo).

**Figure 2.**
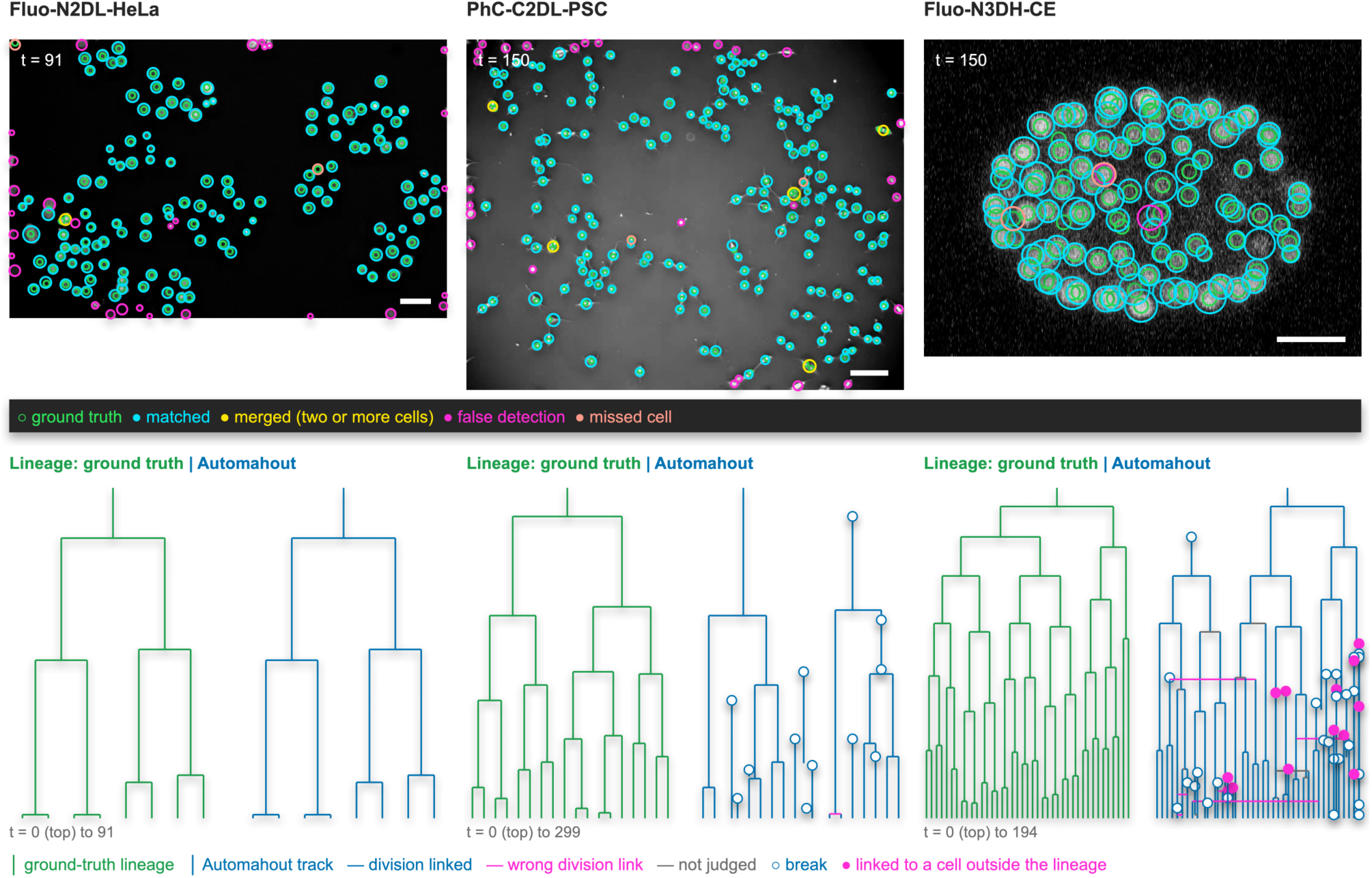
| Autonomous tracking on three datasets. One frame of each dataset’s training sequence 01, Automahout’s final result checked against the ground truth (CTC TRA markers, green circles): matched (cyan), merged (yellow) and false (magenta) detections and missed cells (salmon pink); a detection on a cell but off its centre (on the embryo, by two planes in z) covers less than half of the cell’s marker and is counted both as a missed cell and as a false detection. HeLa t = 91, scale bar 50 µm; PSC t = 150, 100 µm; embryo t = 150, a maximum-intensity projection over z = 14.5–19.5 µm with circles for objects centred within one cell radius (1.7 µm) of it, 10 µm. Grayscale, display range the 1st–99.8th percentile of each image. In the whole frame, HeLa has 137 ground-truth cells (133 matched, 1 merged, 29 false detections, 2 missed), PSC 187 (174, 5, 28, 2) and the embryo 192 (189, 0, 3, 3). Below each, one ground-truth lineage, and the Automahout tracks that hold its cells on the same layout, each at the ground-truth track it holds most. A horizontal line is a division Automahout linked, magenta where the daughter’s first cell does not follow the mother’s last one in the ground truth (a division found a few frames late counts as linked), and grey where the ground truth cannot judge it because the mother’s last cell or the daughter’s first lies on no cell of the ground-truth lineage shown; an open circle is a track that starts without a parent (a break), and a magenta dot a track linked to a cell outside the ground-truth lineage. The lineages shown are those of ground-truth tracks 225 (HeLa, 7 divisions), 881 (PSC, 17) and 640 (embryo, 40); 15, 33 and 107 Automahout tracks hold their cells, with 7, 9 and 22 divisions linked correctly, 0, 2 and 6 wrong division links, 0, 0 and 13 that could not be judged, 0, 0 and 13 links to a cell outside the lineage, and 0, 12 and 27 breaks. Scores, detectors and the operations each dataset needed are in Table 1.

**Table 1.** | Autonomous tracking on three datasets. Scores (on each dataset’s training sequence 01) as defined in Methods.

| Dataset (training sequence 01) | Modality | Detector, and how it was chosen | Operations carried out | DET | TRA | LNK | BC(0) | Time |
|---|---|---|---|---|---|---|---|---|
| Fluo-N2DL-HeLa | 2D fluorescence | Cellpose-SAM: agreed best with the image | pretrained detector as it is; link; lineage | 0.954 | 0.954 | 0.953 | 0.83 | 31 min |
| PhC-C2DL-PSC | 2D phase contrast | Cellpose-SAM: persisted best from frame to frame | pretrained detector as it is; link; lineage | 0.956 | 0.954 | 0.938 | 0.39 | 49 min |
| Fluo-N3DH-CE | 3D fluorescence | ELEPHANT: no pretrained detector persisted | annotate; train from random weights; review and correct; link; lineage | 0.971 | 0.965 | 0.920 | 0.55 | 4.5 h |

Linking is the weaker half. Links from one frame to the next are mostly right (LNK 0.95 on HeLa, 0.94 on PSC, 0.92 on the embryo), but whole tracks break: 61% (HeLa), 19% (PSC) and 25% (embryo) of the ground-truth tracks are recovered complete from start to end (CT), and divisions are recovered less often than cells (BC(0) 0.83, 0.39 and 0.55). Automahout links with ELEPHANT’s nearest-neighbour linker, which has no model of motion: in crowded frames, a cell that moves farther than the distance to its neighbour, or daughters that move apart quickly after a division, break a track, and the lineage passes repair only some of these breaks. ELEPHANT can also learn the cells’ motion from the data with its flow model, trained from annotated links as its detector is from annotated cells [5]. Training it adaptively, window by window, as Automahout already does for the detector, and taking up more refined linking algorithms as they appear, are the next steps.

## Discussion

Automahout shows that the operations an expert carries out to adapt cell tracking to a new dataset, from detection to lineage correction, can be performed by an agent without human input. On three datasets that differ in dimension and modality, it used a pretrained detector where one fitted the data and trained its own from random weights where none did, and produced lineages with TRA scores of 0.95–0.96 against a ground truth it never used.

Other tools automate parts of this work. Generalist detectors such as StarDist and Cellpose remove the need for annotation where they fit the data [1, 2, 18], and human-in-the-loop training in Cellpose and ELEPHANT reduces it where they do not [5, 19]. Tracking frameworks such as TrackMate and Ultrack combine detectors and linkers [20, 21], but the choice of method, its settings and the judgement of its output remain a person’s. LLM agents for bioimage analysis mostly act on request, one step at a time (Omega [9], BioImage.IO Chatbot [12]). More recent multi-agent systems carry out a whole workflow from a request, operating ImageJ (CopilotJ [11]), writing Fiji scripts (Agentic-J [10]) or segmenting and analysing organoids (MorphoOrgaAgent [22]), but with existing models used as they are, not trained and corrected on the data. Automahout differs in closing the loop: it judges its own output against references it made, decides what to correct, and decides when to stop.

This preprint introduces the concept and shows it on three datasets, on one training sequence (01) each and in one run each. Its scores are therefore not comparable with the challenge’s published results, which are on the test sequences. For the full paper we plan a broader evaluation: further Cell Tracking Challenge datasets, their test sequences through a regular challenge submission, repeated runs, which operations matter, and a comparison with an expert using ELEPHANT. Several limits are already clear. Because the reference cells Automahout annotates for itself are bright blobs on a dark background, its review applies to fluorescence; on phase contrast it currently depends on a pretrained detector passing its check. Because some references are not cells, the stopping rule tolerates up to 3% missed references and can therefore stop with a small number of real cells still unlearned. None of its criteria is ground truth, and a result that satisfies all of them can still be wrong. The vision-language model judges large, well-separated cells well and small, crowded ones poorly, which is why its eye is checked on every frame before its answer is used. Even after that check it can be wrong, and on the embryo it rejected a real cell (Supplementary Fig. 1). The error is contained, since a rejected reference is only dropped, never taught as background, but that cell is no longer required of the detector. The check also currently asks whether a reference is a cell nucleus, which suits nuclear labels; asking instead whether it is a cell or the labelled part of one will extend it to other labels. More capable vision-language models, or models trained on microscopy images, should make these judgements more reliable; because the model is a setting, a better one can be used without changing the rest.

The operations are implemented as tools, plugins that declare what data they apply to and can also be called on their own. A new algorithm is therefore taken up by adding a tool, without changing the rest, and Automahout chooses among the candidates for each dataset by its own checks, the right tool for each dataset. Adding Cellpose-SAM as one more candidate detector was such a change: Automahout’s own check chose it on both cultures, and PSC’s TRA rose from 0.89 to 0.95, while on the embryo, where no pretrained detector persisted, it still trained its own. Linking is where we will use this next: a learned linker, such as ELEPHANT’s flow model trained on the run’s own links or a future algorithm, can be added as a tool and chosen where it does better than nearest-neighbour linking. What Automahout shows is that the operations a person carries out to track a new dataset (annotating, training, inferring, evaluating, correcting and deciding when to move on) can be carried out without a person. As detectors, references and models improve, the same operations should carry over, still without a person in them.

---

**Supplementary Table 1.**
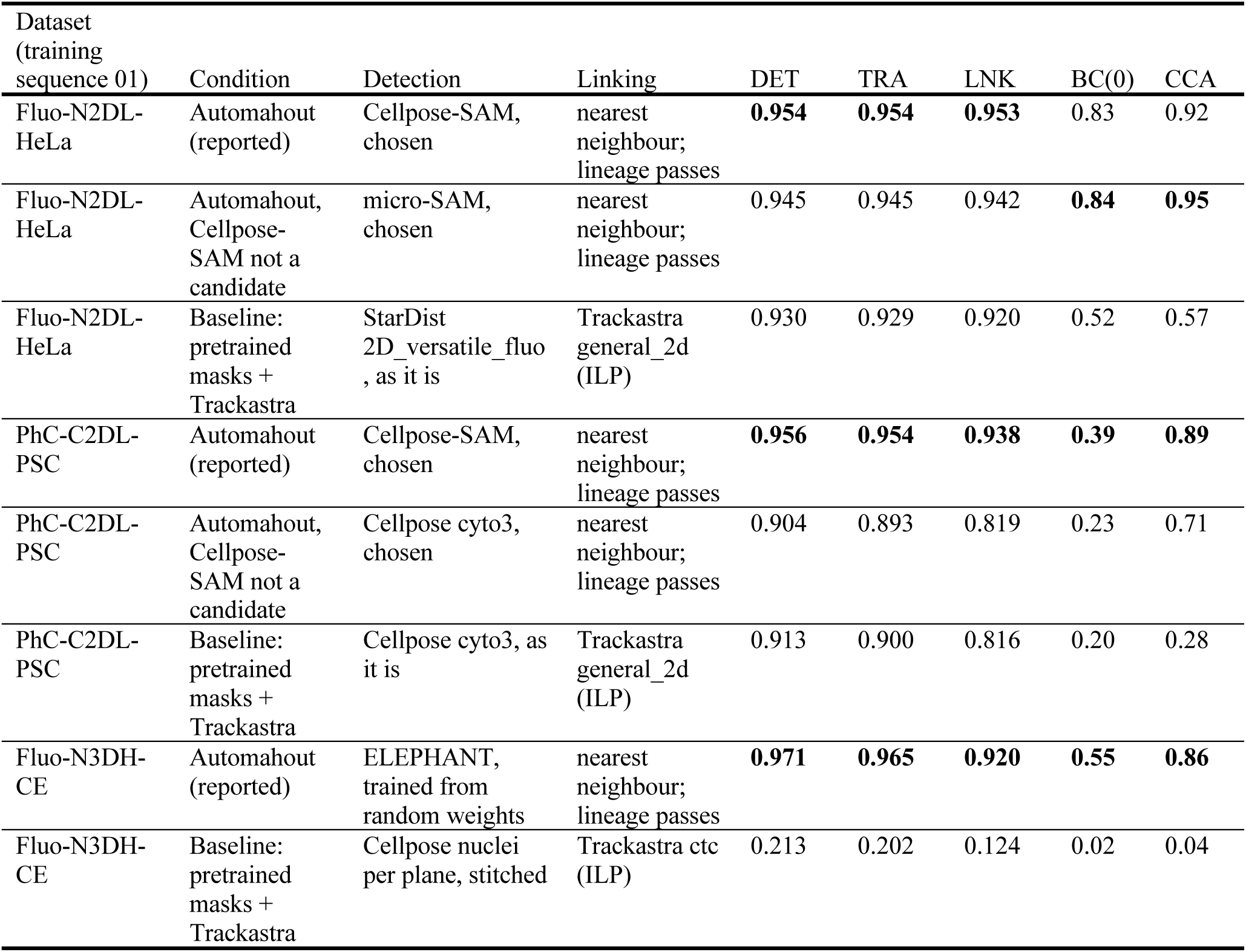
| Other conditions and a baseline. The baseline (scripts/baselines/trackastra_baseline.py) applies one pretrained detector as it is and links its masks with Trackastra 0.5.6 [17] (ILP; model general_2d on the cultures, ctc on the embryo), with no review, correction or lineage passes. On the embryo the masks are the limit: at t = 50, 100 and 150 the masks cover more than half of 0, 16 and 55 of the 26, 93 and 192 ground-truth cells. All scores are on each dataset’s training sequence 01 with py-ctcmetrics 1.3.3 [23]. In each dataset, the best value of each score is in bold. Scores as defined in Methods.

| Dataset (training sequence 01) | Condition | Detection | Linking | DET | TRA | LNK | BC(0) | CCA |
|---|---|---|---|---|---|---|---|---|
| Fluo-N2DL-HeLa | Automahout (reported) | Cellpose-SAM, chosen | nearest neighbour; lineage passes | **0.954** | **0.954** | **0.953** | 0.83 | 0.92 |
| Fluo-N2DL-HeLa | Automahout, Cellpose-SAM not a candidate | micro-SAM, chosen | nearest neighbour; lineage passes | 0.945 | 0.945 | 0.942 | **0.84** | **0.95** |
| Fluo-N2DL-HeLa | Baseline: pretrained masks + Trackastra | StarDist 2D_versatile_fluo, as it is | Trackastra general_2d (ILP) | 0.930 | 0.929 | 0.920 | 0.52 | 0.57 |
| PhC-C2DL-PSC | Automahout (reported) | Cellpose-SAM, chosen | nearest neighbour; lineage passes | **0.956** | **0.954** | **0.938** | **0.39** | **0.89** |
| PhC-C2DL-PSC | Automahout, Cellpose-SAM not a candidate | Cellpose cyto3, chosen | nearest neighbour; lineage passes | 0.904 | 0.893 | 0.819 | 0.23 | 0.71 |
| PhC-C2DL-PSC | Baseline: pretrained masks + Trackastra | Cellpose cyto3, as it is | Trackastra general_2d (ILP) | 0.913 | 0.900 | 0.816 | 0.20 | 0.28 |
| Fluo-N3DH-CE | Automahout (reported) | ELEPHANT, trained from random weights | nearest neighbour; lineage passes | **0.971** | **0.965** | **0.920** | **0.55** | **0.86** |
| Fluo-N3DH-CE | Baseline: pretrained masks + Trackastra | Cellpose nuclei per plane, stitched | Trackastra ctc (ILP) | 0.213 | 0.202 | 0.124 | 0.02 | 0.04 |

**Supplementary Figure 1.**
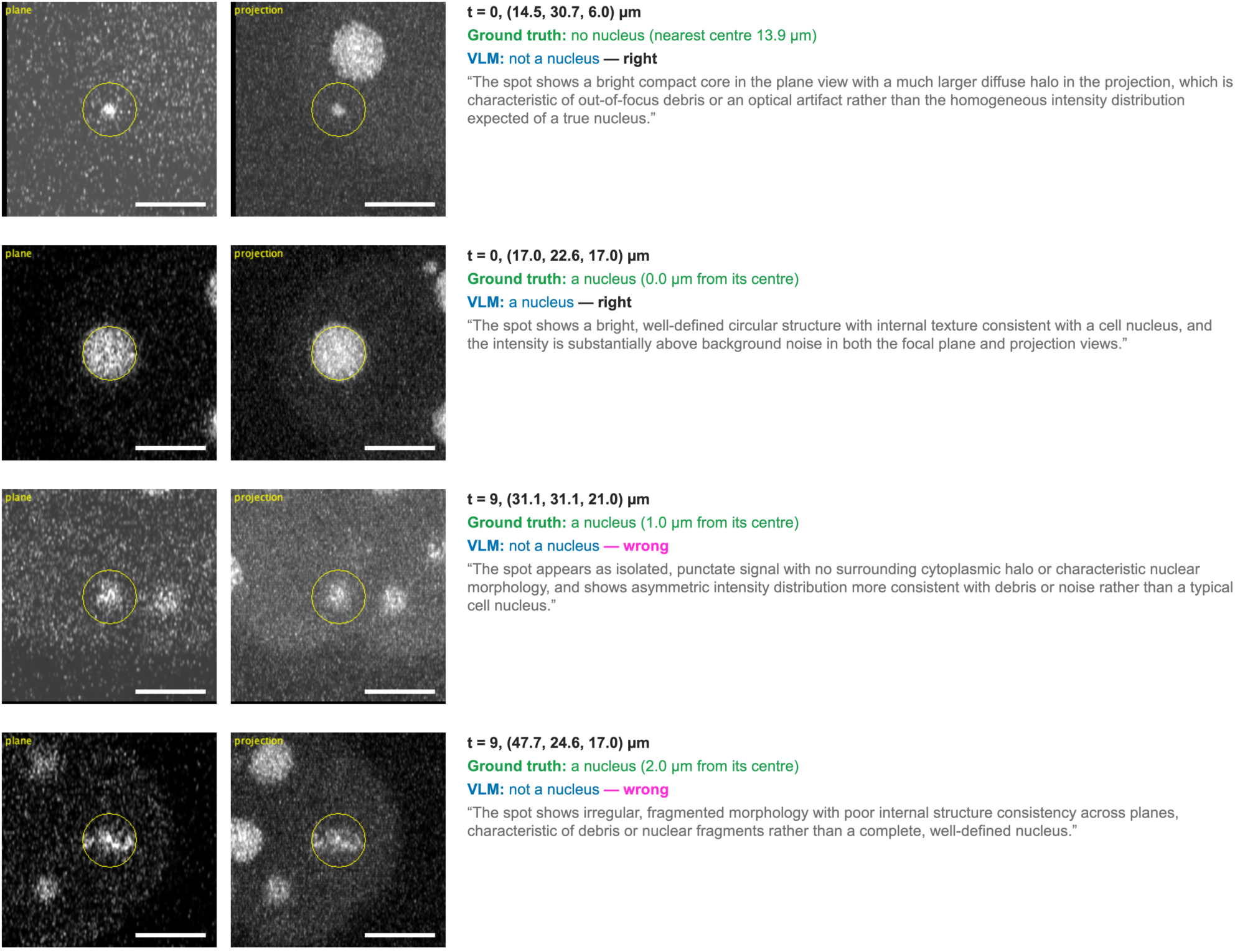
| What the vision-language model was shown and what it answered. What the vision-language model was shown for a doubtful reference on the embryo (left, the plane through it; right, a projection over the planes around it; the circle is the cell radius measured in the window, 3.81 µm; each crop is 30.5 µm across; scale bar, 10 µm), its answer and its reasoning, beside the ground truth (a reference is on a nucleus when a CTC TRA marker centre lies within one radius). It rejects the polar body at t = 0 and accepts a nucleus, but rejects two real nuclei at t = 9; a rejected reference is therefore only dropped, never taught as background. The question was replayed offline on these four references of the first window. In the run reported here it was consulted once, at t = 5, passed the check and rejected one reference, also a real nucleus (0.2 µm from a ground-truth centre), which was dropped.

**Supplementary Figure 2.**
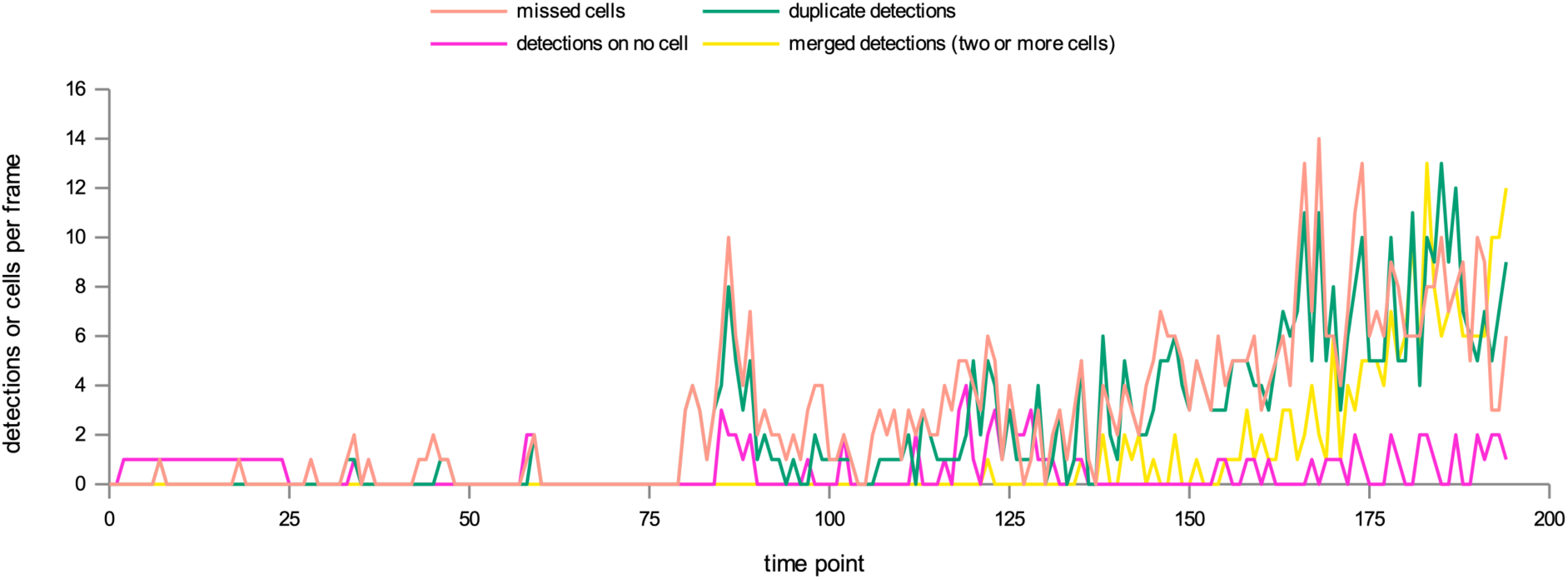
| Errors per frame on the embryo. Missed cells, duplicate detections, detections on no cell and merged detections per frame against the ground truth (CTC TRA markers). They stay low until the embryo crowds, after t ≈ 160.

**Supplementary Figure 3.**
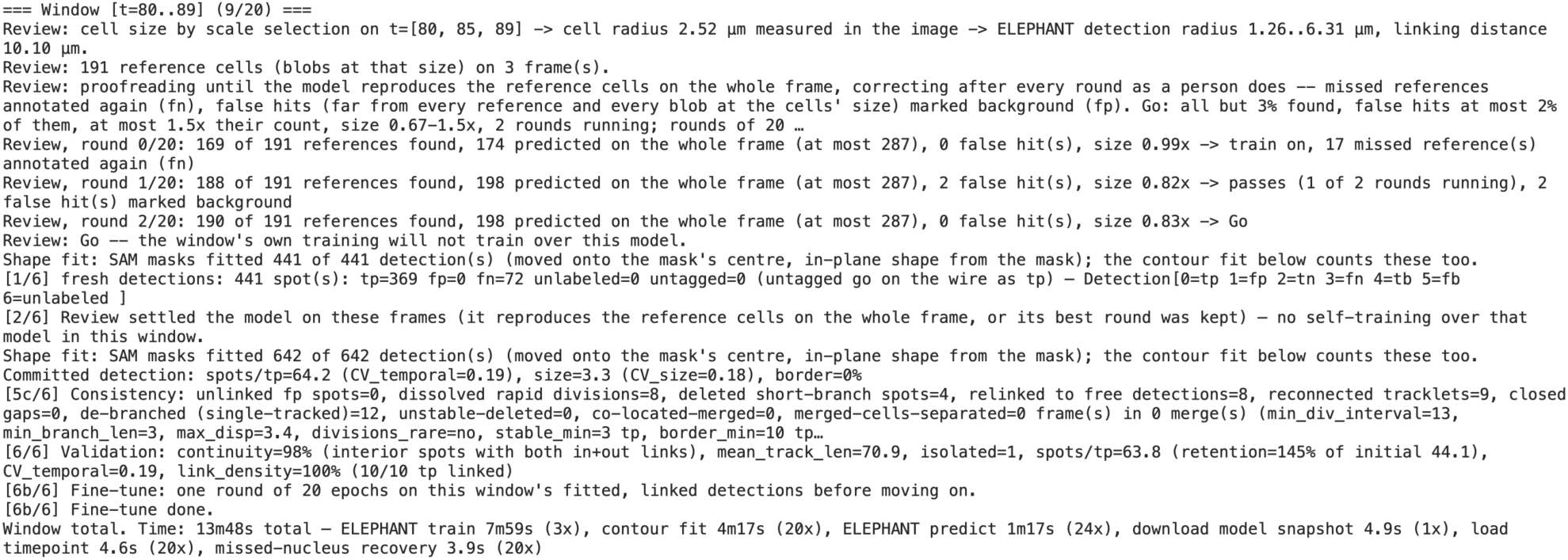
| The run log of one time window. The lines of the embryo run’s log for t = 80–89 a person would read: what Automahout measured, decided and corrected, round by round, without a question to a person.

**Supplementary Figure 4.**
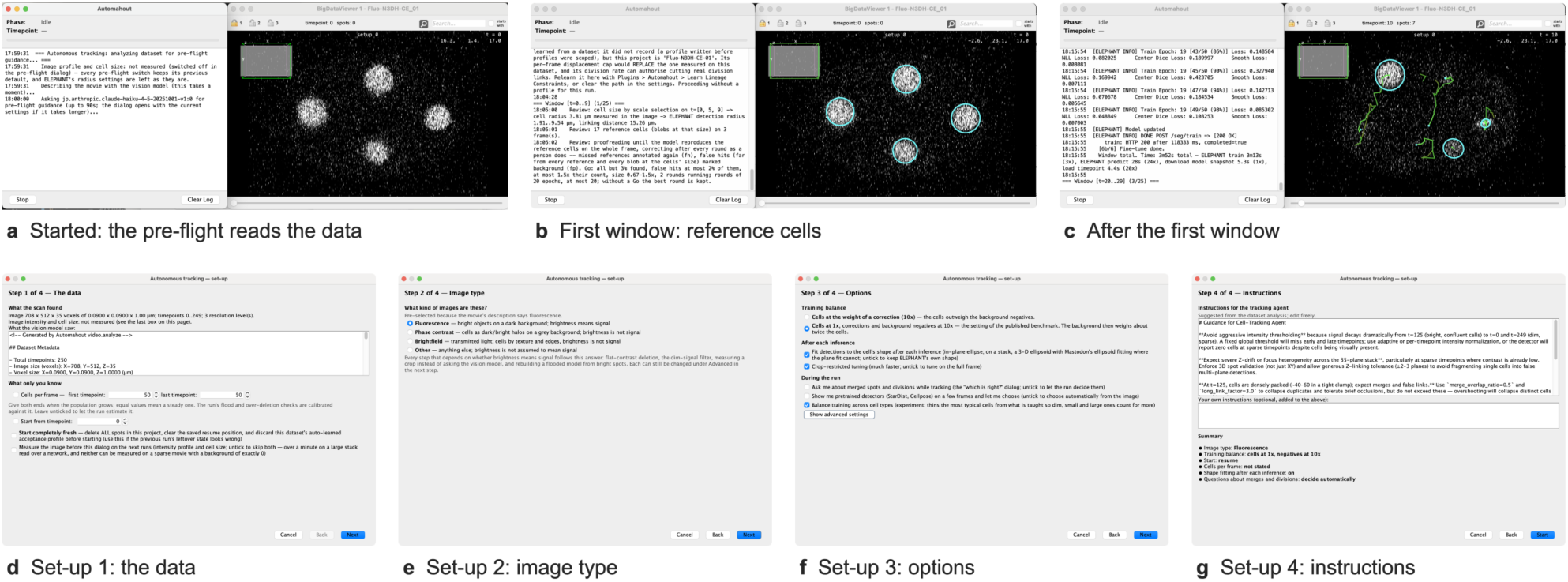
| Automahout in Fiji. A run on the embryo (Fluo-N3DH-CE 01) started from Mastodon’s Plugins > Automahout menu, with Automahout’s log beside BigDataViewer. (**a**) The pre-flight reads the data before the set-up opens. (**b**) The first window’s review: the cell radius measured in the image and the reference cells annotated at it (cyan). (**c**) The end of the first window: the fine-tune on its linked detections, and the tracks (green). (**d**–**g**) The set-up, in four steps: what the scan and the vision-language model found and what only the user knows (d), the image type (e), the training balance and options (f), and the instructions for the agent, suggested from the dataset analysis and editable (g). In the GUI the agent chooses the order of the steps; the runs reported here used the same plugin code with a fixed order and the same training balance (cells at 1x).

## Methods

### Implementation

Automahout is a Java plugin for Mastodon [6] (Fiji [13]). ELEPHANT detects and trains on a server reached over HTTP; pretrained detectors (StarDist [1], Cellpose 3 [16], Cellpose-SAM [4] and micro-SAM [3]) and shape fitting (micro-SAM) run on tool servers with a small HTTP/JSON protocol. Vision questions go to Claude Haiku 4.5 (Anthropic) through Amazon Bedrock. Each operation is a tool that can also be called on its own. Fiji, Mastodon, Automahout and the tool servers ran on a MacBook Pro (Apple M2: 8-core CPU, 10-core GPU; 24 GB RAM; macOS 14.3.1); the ELEPHANT server ran in a Docker container on a server (AMD EPYC 7643, 48 cores; 512 GB RAM; Ubuntu 22.04.5 LTS) with one NVIDIA H100 PCIe GPU (80 GB; driver 590.48.01). Details and measurements are in Supplementary Note 1.

### Sequence

Before the first window, Automahout reads the modality from the intensity distribution and tries the pretrained detectors on frame pairs at the start, middle and end of the movie, keeping one only if at least 80% of its detections persist into the next frame (on 2D fluorescence, if they agree with the bright regions). A kept detector is used as it is. Otherwise an ELEPHANT model is created from random weights. Either way the movie is processed in windows of 10 frames, each detected (with ELEPHANT, after its review), linked and corrected in turn, followed by a lineage correction over the movie.

### Cell size and references

Each window’s cell radius is read from the image by scale selection [24]: the median radius at which scale-normalised Laplacian-of-Gaussian blobs peak, searched within 0.5–2 times the previous window’s radius and 1–10 µm. ELEPHANT’s radius range and linking distance follow it. On fluorescence, the reference cells are the blobs at that radius on three frames of the window whose response is at least half the frame’s median, placed as ELEPHANT annotations.

### Review and stopping

ELEPHANT is trained in rounds of 20 epochs. After each round the whole frame is predicted and compared with the references: found, missed, false hits (detections far from every blob) and size. Missed references are re-tagged as missed cells once; false hits are marked background, within 25% of the annotations. Go requires, in two rounds running, at most 3% of references missed, false hits at most 2%, predictions at most 1.5 times the references and a size ratio of 0.67–1.5; after 20 rounds without Go the best round is kept. After linking and lineage correction, ELEPHANT trains one more round on the window’s detections, shape-fitted by micro-SAM (or an ellipse on the cell’s outline), before the next window, whose review checks that model first. A reference missed repeatedly is shown to the vision-language model, whose answer is used only on frames where it first recognises two references the detector finds.

### Linking and evaluation

Detections are linked by ELEPHANT’s nearest-neighbour linker and corrected per frame (long links relinked, trifurcations resolved) and per window’s lineage (divisions too soon after the last, or whose daughters do not move apart or differ in form, dissolved; short branches deleted; breaks joined across up to 3 frames; tracks shorter than 3 frames deleted; one cell tracked twice merged). Results are exported in the Cell Tracking Challenge format and scored with py-ctcmetrics [23] against the ground truth, which was used only to score, on each dataset’s first training sequence (01). All scores range from 0 to 1, higher being better [14, 15]. DET and TRA are one minus the normalised cost of the edits that would turn the result into the ground truth: adding, removing, splitting or merging detections for DET, and for TRA also adding, removing or changing links. LNK is the linking part of that cost alone. BC(0) is the F1 score of divisions found at the right frame. CCA compares the distribution of cell-cycle lengths with the ground truth’s. CT is the fraction of ground-truth tracks reconstructed completely, from start to end. No manual input was given: no annotation, region of interest, parameter or answer came from a person. The runs used a fixed sequence of steps over the whole movie (t = 0–194 on the embryo, whose ground truth ends there) with the converged-tracks lineage pass (which splits one detection that two tracks ran into) switched off, the same for all three datasets. The code is in the public repository (https://github.com/automahout/mastodon-automahout) at tag `preprint-v1`, the code the embryo was run with. The cultures were run before two fixes that make two lineage passes measure distance in 3-D on a stack (development commit 0d69e63; described in the repository’s `paper/README.md`); the fixes leave single planes unchanged. Versions of the other software are listed in Supplementary Note 1.

## Data availability

All relevant data are available within the paper and on SSBD:repository (https://doi.org/10.24631/ssbd.repos.2026.11.552): the tracking results in the Cell Tracking Challenge format, the Mastodon projects, the run logs and the embryo’s trained ELEPHANT model. The input images and ground truth are the Cell Tracking Challenge’s datasets Fluo-N2DL-HeLa, PhC-C2DL-PSC and Fluo-N3DH-CE [14, 15], available from the challenge (Supplementary Note 1).

## Acknowledgements

We thank the Cell Tracking Challenge (celltrackingchallenge.net) and its organisers for the datasets used in this work [14, 15]. We thank all members of the Onami laboratory for helpful discussions and technical support.

## Author contributions

K.S.: Conceptualization, Methodology, Software Development, Writing – Original Draft; S.O.: upervision, Writing – Review & Editing.

## Use of generative AI tools

Automahout uses a vision-language model (Claude Haiku 4.5, Anthropic) as one of its components; its role is described in Methods and Supplementary Note 1. Separately, the authors used Claude Code (Claude Opus 5.5, Anthropic) as an assistant to write and test the software and the analysis and figure scripts, and to draft and edit the manuscript. The authors reviewed all text, and checked the code by running it and testing its results; the reported results were produced by the code in the public repository and scored against the Cell Tracking Challenge ground truth. The authors take full responsibility for the content of this manuscript.

## Supplementary Note 1 | Detailed methods

### Implementation

Automahout is a Java plugin for Mastodon (Fiji). Detection and training run on an ELEPHANT server over HTTP (`/seg/train`, `/seg/predict`, `/seg/reset`). Pretrained detectors and shape fitting run on separate tool servers speaking a small HTTP/JSON protocol (automahout-remote/1): StarDist, Cellpose 3, Cellpose-SAM and micro-SAM for detection, micro-SAM for shape fitting. The language models are reached through a single backend interface. The runs reported here used Claude Haiku 4.5 on Amazon Bedrock for vision. Every operation is a tool that can also be called on its own, for example `review.frame`, which reviews any frame’s spots against its reference cells and changes nothing.

### Sequence of operations

The benchmark runs used a fixed sequence: per window of 10 frames, review and proofreading (with ELEPHANT only), then detection, linking and correction; then a global lineage correction. Before the first window: - the image profile reads the modality from the frame’s intensity distribution; - the detector trial runs; - ELEPHANT’s detection model is created from random weights (not from ELEPHANT’s pretrained model, whose training data include Cell Tracking Challenge sequences).

### Choosing a pretrained detector

Each applicable pretrained detector is run on frame pairs (t, t+1) at the first, middle and last frame. - On 2D fluorescence, a detector is kept when its detections agree with the frame’s bright regions (F1 ≥ 0.90). - Otherwise, a detector is kept when at least 80% of its detections persist into the next frame (one-to-one within one cell radius) on every pair.

A kept detector detects the whole movie as it is; its output is linked and corrected but ELEPHANT is not trained. When none is kept, ELEPHANT detects. On data that do not read as fluorescence, a detector that persists on every pair without reaching 80% may still seed ELEPHANT’s first training; this did not arise in these runs.

### Cell size

The cell radius is read per window by scale selection, without the truth or the vision model: 1. The scale-normalised Laplacian-of-Gaussian response is computed at radii 1.25 times apart. 2. The blobs at every radius (local maxima within one radius, above 8 median absolute deviations of the response) are pooled, one per place. 3. Each blob takes the radius at which its response peaks, and the median is the window’s radius.

The search spans 0.5–2 times the previous window’s radius, always within 1–10 µm (the whole of it for the first window); without that bound, a measurement on the lowest rung fed the next window’s search and the radius collapsed late in the embryo. ELEPHANT’s detection radius range (0.5–2.5 times the radius) and linking distance (4 times) are re-centred on it every window, so that predictions of the current cells’ size are not discarded. Measured on Fluo-N3DH-CE 01, every 30th frame: estimated 4.58, 3.66, 2.34, 1.88, 1.88, 1.88 and 1.50 µm against ground-truth median radii of 4.32, 3.55, 2.54, 2.57, 2.38, 2.28 and 1.92 µm. Blobs at the estimate found 94–100% of the ground-truth cells, with at most 4.6% of blobs on no cell.

### Reference cells and review

On fluorescence data, a window’s reference cells are the confident blobs at its cell size on its first, middle and last frame: those whose response is at least half the frame’s median blob response, as a person annotates only cells they are sure of. On Fluo-N3DH-CE 01 t = 0–19 this took the blobs on no cell from 46 of 184 to 9 of 142, for 134 instead of 139 of 140 cells covered. They are placed as ELEPHANT annotations of their radius. A review (`FrameReview`) compares a prediction of the whole frame with the references: - a reference is **found** when a detection’s centre lies within one radius of it, and **missed** otherwise; - a detection is a **false hit** when it lies more than two radii from every reference and more than one from every blob (a blob that is not a reference may be a cell); - the **size ratio** is the median radius of the detections on references over the cell radius.

### Training, proofreading and stopping

ELEPHANT is trained on the window’s annotations in rounds of 20 epochs (5 crops of 256 × 256 (× 16) voxels per frame, learning rate 10⁻³, class weights 1:10:10, correction weight 10). After each round the whole frame is predicted and reviewed, and the review’s findings are fed back as ELEPHANT’s own labels: - a missed reference’s annotation is re-tagged as a missed cell (fn) once; - false hits are marked background (fp) while background stays within 25% of the frame’s annotations; - at most 10% of the references of each kind are corrected per round.

**Go** is declared when, in two rounds running: - all but 3% of the references are found; - false hits are at most 2% of them; - predictions are at most 1.5 times their number; - the size ratio is 0.67–1.5.

Before any training, the current model is reviewed once; a model that already passes is kept unchanged. After 20 rounds without Go, the round with the most references found (false hits and flooding counted against it) is restored. A window settled this way is not self-trained during detection; once it is linked and corrected, ELEPHANT trains one round (20 epochs) on its detections with their fitted shapes, and the next window’s review checks that model before any training. The references themselves stay spheres of the cell radius: fitting them before training (micro-SAM on the plane through the centre, depth kept) stalled the first embryo window, with 11– 14 of 15 references found over 20 rounds, against Go after 2 rounds with spheres.

### Doubtful references

When only missed references keep a round from passing, a reference missed in two rounds running is shown to the vision-language model (at most five per window). It sees the plane through the reference and a projection over ±2 radii in z, eight radii across, with a circle of the cell radius, and is asked whether it is a nucleus. - Its answer is used only on frames where it first calls nuclei two references that the detector finds. - A reference it rejects is dropped from the references, and the detector is no longer required to find it. It is not marked background, because the model can be wrong even after passing the check. In the run reported here it passed the check at t = 5 and rejected one reference, a real cell (0.2 µm from a ground-truth centre); asked offline about references of the first window (Supplementary Fig. 1), it also rejected two real cells at t = 9.

Measured on Fluo-N3DH-CE 01: at t = 0 it accepted the four cells and rejected a stationary bright object. At t = 60 and 180 it rejected seven of seven cells, which the check of its eye prevents from mattering.

### Detection, correction and shape

After the review, each frame of the window is detected on the whole frame and cleaned by the detection passes. These remove duplicates, detections with no signal and malformed ones. On stacks they also re-place one-plane detections in z, and on fluorescence they mark bright cells that no detection covers as missed. Shapes are fitted by micro-SAM (vit_b_lm) point prompts at each detection, falling back to an ellipse fitted to the cell’s contour.

### Linking and lineage correction

Detections are linked by ELEPHANT’s nearest-neighbour linker within the window, with the linking distance from the cell size. Correction passes then run in this order (from the run’s log, `Correction` `plan`): - **on each frame ELEPHANT detects:** extent, split merged, duplicates, flat contrast, no signal, debris, protected overlap, weak, minimum size, one-plane z refit, recover missed, contour fit; - **per frame:** weak and small spots, sandwiched and stacked duplicates, merge duplicates, long links, relink, convergent links, trifurcations, recover predecessors, division daughters, orphan starts, short tracklets, fill gaps, isolated-spot rescue, bridge gaps, isolated spots, anisotropic refit, bridge after refit, isolated (final); - **per window’s lineage:** strip links from background labels; converged tracks (switched off in these runs); measure the caps (per-frame displacement, branch length, division interval); dissolve divisions sooner than the minimum interval; delete short post-division branches; move links onto the cell’s own free detection; join a track end to a start in the next frame within 3 radii, then up to 3 frames later; dissolve divisions whose daughters do not move apart or that fail the interval or the daughters’ form; delete tracks shorter than 3 frames (10 at the image edge); merge two tracks that hold a steady sub-cell offset (one cell tracked twice).

### Evaluation

Results are exported in the Cell Tracking Challenge format and scored with py-ctcmetrics 1.3.3 (DET, TRA, LNK, BC, CCA) against the challenge’s ground truth. Sampled-frame detection uses the CTC DET formula on the listed frames. The truth was used only to score. All three sequences are the challenge’s first training sequence (01), which was also used while developing the method; the scores are therefore not a held-out evaluation and are not comparable with the challenge’s test-set results.

### Software versions

ELEPHANT server: v0.7.1 (elephant-track/elephant-server, tag v0.7.1).

Automahout: the public repository (https://github.com/automahout/mastodon-automahout) at tag `preprint-v1`. Fluo-N3DH-CE 01 was run at development commit dcd7f71, whose pipeline source (`src/main`) is the tag’s but for a development tool the runs do not use. Fluo-N2DL-HeLa 01 and PhC-C2DL-PSC 01 were run earlier, at development commit 0d69e63: its pipeline source differs from the tag’s only by two fixes, which leave single planes unchanged (described in the repository’s `paper/README.md`). Software used in the runs:

| Component | Version |
|---|---|
| Automahout | preprint-v1 (the runs: development commits 0d69e63 and dcd7f71); Java 21.0.4; pom-scijava 42.0.0 |
| Mastodon | 1.0.0-beta-35 (BigDataViewer core 10.6.4, ImgLib2 7.1.5); mastodon-ellipsoid-fitting 0.7.2 |
| ELEPHANT client | 0.8.0 |
| ELEPHANT server | v0.7.1 (tag v0.7.1; the runs used its code at e2ad410, before the release), run as the Docker image built from it (base pytorch/pytorch:2.3.1-cuda12.1-cudnn8-runtime) on one NVIDIA H100 PCIe |
| Detector server | Python 3.10; Cellpose 3.0.10 (cyto3, nuclei; nuclei in 3D); StarDist 0.9.1 with CSBDeep 0.8.0 and TensorFlow 2.16.2 (2D_versatile_fluo, 2D_versatile_he); PyTorch 2.4.0 |
| Cellpose-SAM server | Python 3.12; Cellpose 4.2.1.1 (model cpsam), PyTorch 2.14.1, on Apple M2 (MPS); examples/cellpose-sam |
| Shape-fit server | Python 3.12; micro-SAM 1.8.14 (git 323bef4; model vit_b_lm), Segment Anything 1.0, PyTorch 2.14.1, on Apple M2 (MPS), macOS 14.3.1 |
| Vision-language model | Claude Haiku 4.5 on Amazon Bedrock (jp.anthropic.claude-haiku-4-5-20251001-v1:0) |
| Scoring | py-ctcmetrics 1.3.3 |

Python versions are pinned by each tool’s uv.lock in the repository at the tag.

### Commands

The tool servers, each from its folder in the repository (macOS, Apple M2):

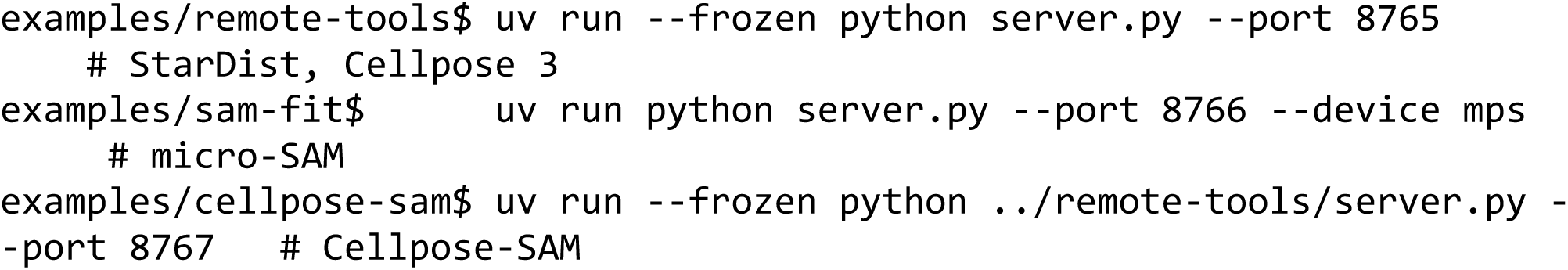

The three runs, from the repository root (`run_drift.sh` builds the named commit, runs Automahout as a batch job with its own settings, without the set-up dialog or any interaction, and scores the result). The ELEPHANT server URL is given as `ELEPHANT_URL`; the vision-language model’s credentials come from the user’s Automahout settings and are not recorded.

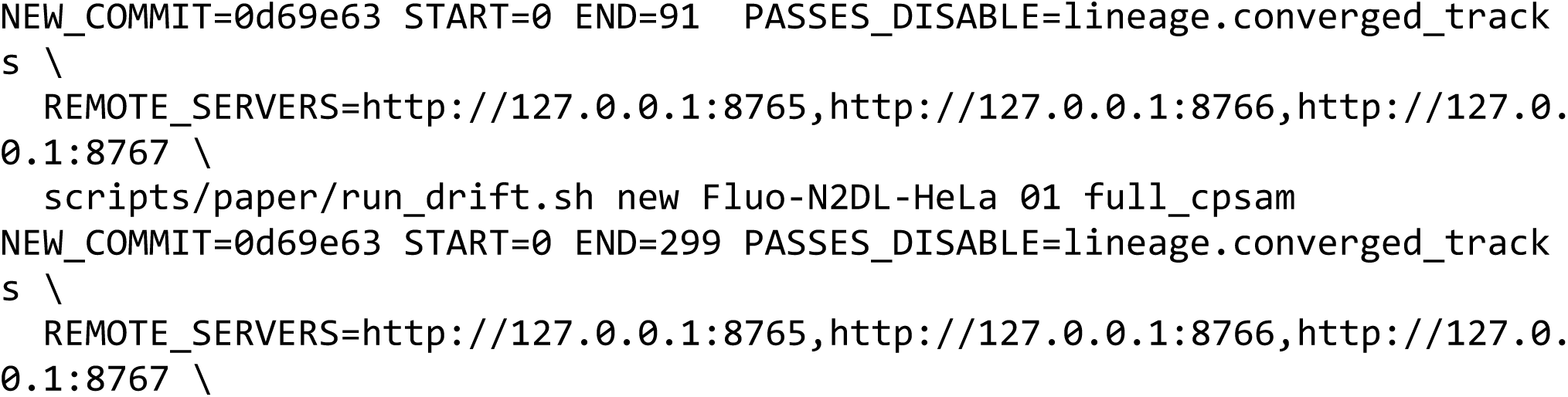
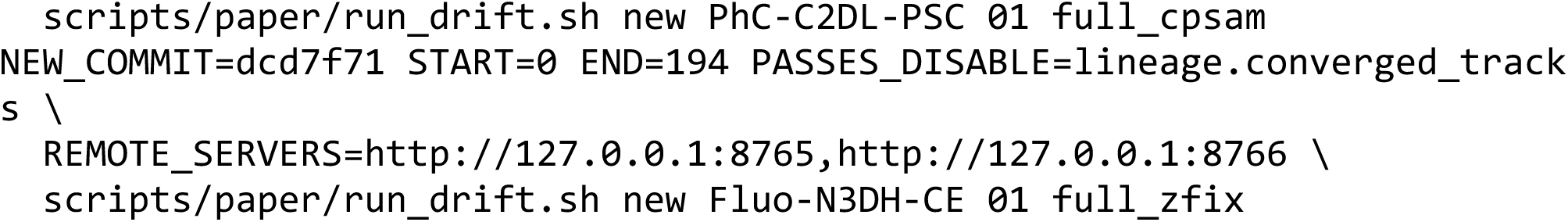

`NEW_COMMIT` names the development commit a run used; as the public repository does not carry those commits, omit `NEW_COMMIT` there to build the version checked out (`preprint-v1`). START and END run the fixed sequence of steps over that range (all of HeLa and PSC; the embryo’s frames with ground truth). Each run writes `run-info.json` (settings, servers, disabled passes), `run.log`, `metrics.json`, `frames.csv`, `windows.json`, the project and `RES/` to `$BENCH_OUT/drift/<dataset>/01/new/rep<name>`.

## Code and data availability

The datasets are the Cell Tracking Challenge’s training datasets Fluo-N2DL-HeLa, PhC-C2DL-PSC and Fluo-N3DH-CE [14, 15], to be downloaded from the challenge (https://celltrackingchallenge.net/2d-datasets/ and https://celltrackingchallenge.net/3d-datasets/); they are not redistributed with this work.

Code: https://github.com/automahout/mastodon-automahout, tag `preprint-v1`, with the figure data and scripts (`paper/figures`, `scripts/paper`). Run outputs (the results in the Cell Tracking Challenge format, the Mastodon projects, the run logs) and the embryo’s trained ELEPHANT model: SSBD:repository (https://doi.org/10.24631/ssbd.repos.2026.11.552).

